# Ubiquitin Links Smoothened to Intraflagellar Transport to Regulate Hedgehog Signaling

**DOI:** 10.1101/2019.12.18.880799

**Authors:** Paurav B. Desai, Michael W. Stuck, Bo Lv, Gregory J. Pazour

**Affiliations:** Program in Molecular Medicine, University of Massachusetts Medical School, Biotech II, Suite 213, 373 Plantation Street, Worcester, MA 01605

**Keywords:** intraflagellar transport, Hedgehog signaling, cilia, ubiquitin, smoothened

## Abstract

In the absence of hedgehog ligand, patched-1 (Ptch1) localizes to cilia and prevents ciliary accumulation and activation of smoothened (Smo). Upon ligand binding, Ptch1 is removed from cilia, Smo is derepressed and accumulates in cilia where it activates signaling. The mechanisms regulating these dynamic movements are not well understood but defects in intraflagellar transport components including Ift27 and the BBSome cause Smo to accumulate in cilia without pathway activation. We find that in the absence of ligand-induced pathway activation, Smo is ubiquitinated and removed from cilia, and this process is dependent on Ift27 and BBSome components. Activation of hedgehog signaling decreases Smo ubiquitination, and ciliary removal, resulting in its accumulation. Blocking ubiquitination of Smo by an E1 ligase inhibitor or by mutating two lysine residues in intracellular loop three cause Smo to aberrantly accumulate in cilia without pathway activation. These data provide a mechanism to control Smo’s ciliary level during hedgehog signaling by regulating the ubiquitination state of the receptor.

**Summary:** Hedgehog signaling involves the dynamic movement of receptors and effectors in and out of cilia. We find that the dynamics of Smo is regulated by ubiquitination, which regulates its interaction with the intraflagellar transport system to control ciliary levels of this receptor.

## Introduction

Primary cilia play critical roles in development by monitoring the extracellular environment and transmitting this information to the cell body and nucleus, allowing the cell to coordinate its physiology with surrounding cells. In mammals and other vertebrates, ciliary dysfunction leads to a variety of structural birth defects in the brain, heart, lungs, kidneys and other organs along with craniofacial and other skeletal abnormalities. While numerous receptors are localized in cilia, hedgehog signaling is the best studied ciliary pathway. This pathway plays fundamental roles during development and many of the developmental defects caused by ciliary dysfunction can be attributed to abnormal hedgehog signaling. All the key components of the hedgehog pathway are enriched in the cilium (Corbit et al., 2005; Haycraft et al., 2005; Ocbina and Anderson, 2008; Rohatgi et al., 2007) and their localization changes dynamically in response to the activity of the pathway. In the off-state, patched-1 (Ptch1), the hedgehog ligand receptor, accumulates in cilia and prevents ciliary accumulation and activation of smoothened (Smo). Upon binding of ligand, Ptch1 is removed from the cilium, Smo is derepressed and now accumulates in the cilium. Smo subsequently activates downstream signaling, which results in the accumulation of the Gli transcription factors at the ciliary tip before their modification and translocation to the nucleus where they modulate expression of target genes.

The mechanisms underlying the trafficking of hedgehog components to and within the cilium are not well understood. Part of the movement is facilitated by intraflagellar transport (IFT) (Eguether et al., 2018; Eguether et al., 2014; Keady et al., 2012) and perturbing IFT disrupts hedgehog signaling (Huangfu et al., 2003; Liem et al., 2012). IFT, which is critical for assembly and maintenance of cilia, involves motor driven transport of large complexes called IFT particles along the cilia. These particles are composed of at least 30 proteins organized in IFT-A, IFT-B and BBSome subcomplexes. The IFT particles serve as motor adaptors connecting various proteins that need to be moved into or out of cilia to kinesin and dynein motors. We previously showed that Ift25 and Ift27, which are subunits of IFT-B, are not required for ciliary assembly. Instead, these two IFTs work with the adaptor protein Lztfl1 and the BBSome to regulate hedgehog signaling and maintain proper levels of Smo and Ptch1 in cilia during signaling (Eguether et al., 2018; Eguether et al., 2014; Keady et al., 2012).

In this work, we explore the mechanism underlying the dynamics of Smo localization to cilia and find that it is regulated by ubiquitination. Ubiquitination involves the covalent attachment of the 76 amino acid peptide ubiquitin (Ub) to cellular proteins. Ub is usually added to lysine residues and it can be further ubiquitinated on one of its seven lysine residues to produce polyubiquitinated proteins. The type of Ub modification on a protein determines its cellular fate. For instance, K48 polyubiquitination drives degradation by the proteasome, K11 polyubiquitination drives degradation during specific points in the cell cycle and K63 polyubiquitination often regulates complex formation and signaling (Swatek and Komander, 2016). The addition of Ub results from a cascade of activity where an E1 enzyme activates Ub, passes it to an E2 enzyme, which then either passes the Ub to an E3 enzyme for attachment to the protein of interest or works with an E3 to modify that protein. The process is reversable by the action of deubiquitinating enzymes (Swatek and Komander, 2016). Prior work identified components of the Ub system in cilia (Huang et al., 2009; Pazour et al., 2005; Raman et al., 2015) where it has been implicated in disassembly (Huang et al., 2009; Wang et al., 2019). The Ub system is also known to regulate a number of steps in hedgehog signaling (Hsia et al., 2015). In this work, we find that ubiquitination of Smo facilitates its interaction with the IFT system for efficient ciliary removal/export when the hedgehog pathway is suppressed

## Experimental Procedures

### Plasmids

Plasmids were assembled by Gibson assembly (NEB, Ipswitch MA) into the pHAGE lentiviral backbone (Wilson et al., 2008). All inserts are derived from mouse unless otherwise stated. Mutations were generated by PCR amplification with mutated primers and the products Gibson assembled. To mutate the 16 lysines in the C-terminal tail of Smo, the tail sequence was chemically synthesized (gBlock, IDT, Skokie Illinois) and PCR amplified for Gibson assembly. All inserts were fully sequenced and matched NCBI reference sequence or expected mutant forms. Plasmids are described in Table 1 and SnapGene files will be provided upon request.

**Table 1.** Plasmids.

| Name | Description | Promoter | Insert | Tags | Selection |
| --- | --- | --- | --- | --- | --- |
| MS03 | SHH Expression Plasmid | CMV | HsSHH | none | Bsd |
| GP734 | Flag-Ub | CMV | Ub | Flag, Ub | Bsd |
| PD19 | Flag-Ub<sup>noK</sup> (7 Lys mutated to Arg) | CMV | Ub<sup>noK</sup> | Flag, Ub<sup>noK</sup> | Bsd |
| PD14 | HA-Ub | CMV | Ub | HA, Ub | Puro |
| PD13 | HA-Ub<sup>noK</sup> (7 Lys mutated to Arg) | CMV | Ub<sup>noK</sup> | HA, Ub<sup>noK</sup> | Puro |
| PD22 | Smo-Flag | CMV | Smo | Flag | Bsd |
| GP736 | Smo-Flag-Ub | CMV | Smo | Flag, Ub | Bsd |
| GP777 | Smo<sup>noK</sup>-Flag | CMV | Smo<sup>noK</sup> | Flag | Bsd |
| GP806 | Smo<sup>noK</sup>-Flag-Ub | CMV | Smo<sup>noK</sup> | Flag, Ub | Bsd |
| GP807 | Smo<sup>noK</sup>-Flag-Ub<sup>noK</sup> | CMV | Smo<sup>noK</sup> | Flag, Ub<sup>noK</sup> | Bsd |
| PD23 | SmoM2-Flag | CMV | Smo<sup>W539L</sup> | Flag | Bsd |
| GP735 | SmoM2-Flag-Ub | CMV | Smo<sup>W539L</sup> | Flag, Ub | Bsd |
| GP938 | SmoPi-Flag | CMV | Smo<sup>R455A</sup> | Flag | Bsd |
| GP940 | SmoPi-Flag-Ub | CMV | Smo<sup>R455A</sup> | Flag, Ub | Bsd |
| GP757 | Sstr3-Flag | CMV | Sstr3 | Flag | Bsd |
| GP758 | Sstr3-Flag-Ub | CMV | Sstr3 | Flag, Ub | Bsd |
| GP799 | Smo-Flag | GgCryD1 | Smo | Flag | Bsd |
| GP805 | Like GP799 but with the 21 cytoplasmic lysines of Smo mutated to arginine |  |  |  |  |
| GP802 | Like GP799 but with the two lysines in loop two of Smo mutated to arginine |  |  |  |  |
| GP803 | Like GP799 but with the three lysines (KKK to RRR) in loop three of Smo mutated to arginine |  |  |  |  |
| GP804 | Like GP799 but with the 16 lysines in C-terminal tail of Smo mutated to arginine |  |  |  |  |
| GP824 | Like GP799 but with two lysines (KKK to RRK) in loop three of Smo mutated to arginine |  |  |  |  |
| GP825 | Like GP799 but with two lysines (KKK to RKR) in loop three of Smo mutated to arginine |  |  |  |  |
| GP826 | Like GP799 but with two lysines (KKK to KRR) in loop three of Smo mutated to arginine |  |  |  |  |
| GP821 | Like GP799 but with one lysine (KKK to RKK) in loop three of Smo mutated to arginine |  |  |  |  |
| GP822 | Like GP799 but with one lysine (KKK to KRK) in loop three of Smo mutated to arginine |  |  |  |  |
| GP823 | Like GP799 but with one lysine (KKK to KKR) in loop three of Smo mutated to arginine |  |  |  |  |

### Cell Culture

Wild type, *Ift27^-/-^* and *Bbs2^-/-^* mouse embryonic fibroblasts (MEFs) were derived from E14 embryos and immortalized with SV40 Large T antigen. *Barr1^-/-^/Barr2^-/-^* double knockout MEFs were obtained from R. Lefkowitz (Duke University) and immortalized with SV40 Large T antigen. *Lztfl1^-/-^* cells (Figure S2) were obtained by genome editing of immortalized wild type MEFs using guide (gMS04: GCTCGATCAAGAAAACCAAC) cloned into pLentiCrisprV2 (Addgene plasmid # 52961) (Sanjana et al., 2014). All MEFs were cultured in 95% DMEM (4.5 g/L glucose), 5% fetal bovine serum, 100 U/ml penicillin, and 100 microgram/ml streptomycin (all from Gibco-Invitrogen).

IMCD3 cells (Rauchman et al., 1993) were cultured in 47.5% DMEM (4.5 g/L glucose), 47.5% F12, 5% fetal bovine serum, 100 U/ml penicillin, and 100 microgram/ml streptomycin (all from Gibco-Invitrogen). IMCD3*^Ift27-/-^* cells were obtained from Max Nachury (University of California, San Francisco).

For SAG experiments, MEFs were plated at near confluent densities and serum starved (same culture medium described above but with 0.25% FBS) for 48 hr prior to treatment to allow ciliation. SAG (Calbiochem) was used at 400 nM.

SHH conditioned medium was generated from HEK cells transfected with a SHH expression construct (MS03). Cells stably expressing MS03 were grown to confluency in 90% DMEM (4.5 g/L glucose), 10% fetal bovine serum, 100 U/ml penicillin, and 100 microgram/ml streptomycin, medium was then replaced with low serum medium (0.25% fetal bovine serum) and grown for 48 hrs. Medium was collected, filter sterilized and titered for ability to relocated Smo to cilia. Dilutions similar in effect to 400 nM SAG were used for experiments.

### Lentivirus Production

Lentiviral packaged pHAGE-derived plasmids (Wilson et al., 2008) were used for transfection. These vectors were packaged by a third-generation system comprising four distinct packaging vectors (Tat, Rev, Gag/Pol, and VSV-g) using HEK 293T cells as the host. DNA (Backbone: 5 μg; Tat: 0.5 μg; Rev: 0.5 μg; Gag/Pol: 0.5 μg; VSV-g/: 1 μg) was delivered to the HEK cells using as calcium phosphate precipitates. After 48 hrs, supernatant was harvested, filtered through a 0.45-micron filter and added to subconfluent cells. After 24 hrs, cells were selected with puromycin (Puro, 1 microgram/ml) or blasticidin (Bsd, 60 micrograms/ml for CMV promoter or 20 micrograms/ml for GgCryD1 promoter).

### Immunofluorescence

Cells were fixed with 2% paraformaldehyde for 15 min, permeabilized with 0.1% Triton-X-100 for 2 min and stained as described (Follit et al., 2006). In some cases, fixed cells were treated with 0.05% SDS for 5 min before prehybridization to retrieve antigens. The primary antibodies are described in Table 2.

**Table 2.**
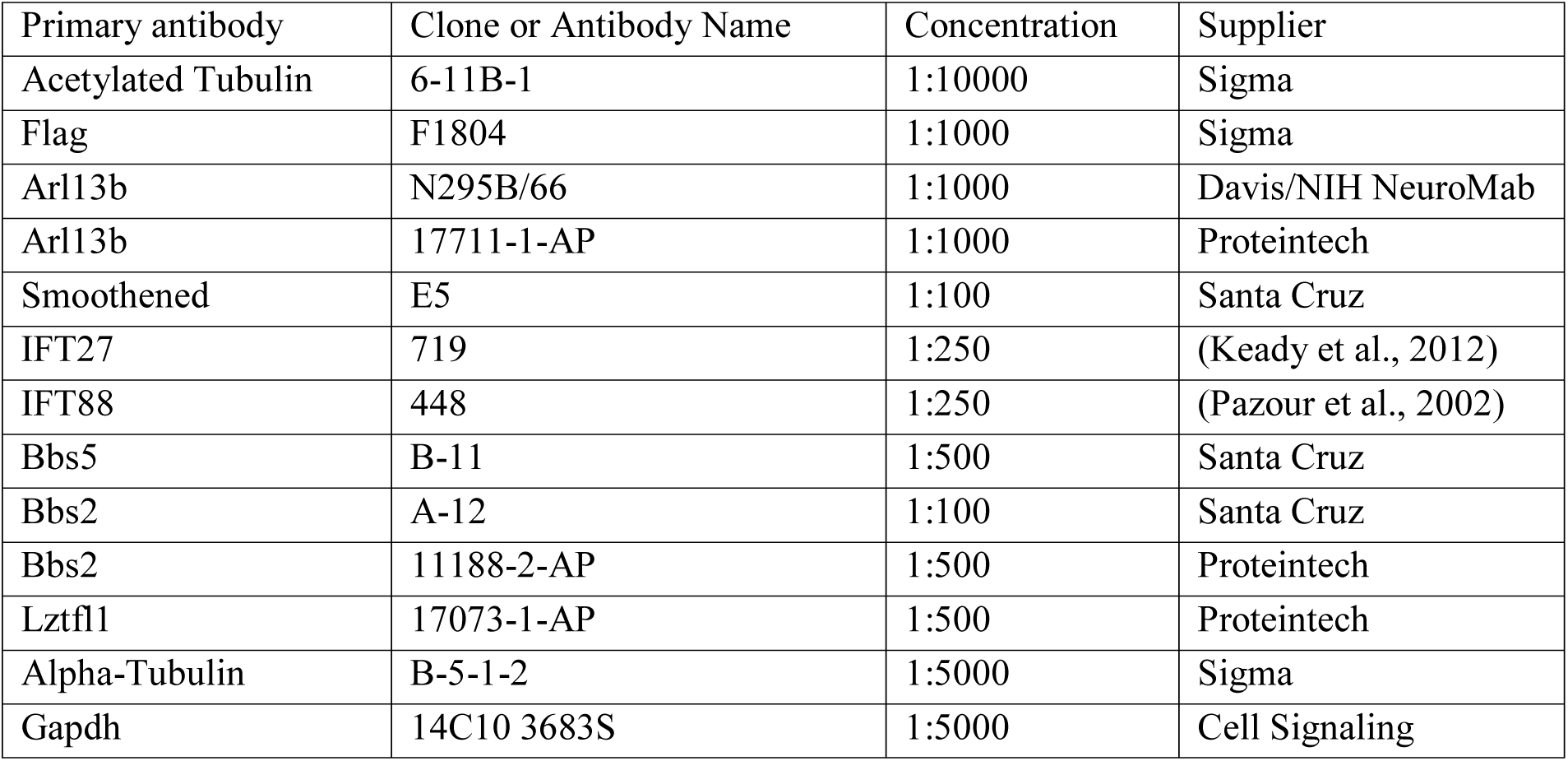
Antibodies.

Confocal images were acquired with an inverted microscope (TE-2000E2; Nikon) equipped with a Solamere Technology – modified spinning disk confocal scan head (CSU10; Yokogawa). Three image Z stacks were acquired at 0.2-micron intervals and converted to single planes by maximum projection with MetaMorph software (MDS Analytical Technologies). Widefield images were captured using an Orca ER or an FLIR camera on a Zeiss Axiovert 200M microscope equipped with a 100x Zeiss objective (Thornwood, NY). Contrast adjustment and cropping was done in ImageJ or Photoshop.

### Protein and mRNA Analysis

For western blots, MEFs were lysed directly into denaturing gel loading buffer (Tris-HCl 125 mM pH6.8, glycerol 20% v/v, SDS 4% v/v, β-mercaptoethanol 10% v/v, bromophenol blue). Western blots were developed by chemiluminescence (Super Signal West Dura, Pierce Thermo) and imaged using an Amersham Imager 600 or Biorad ChemiDoc XRS+.

For immunoprecipitations, cells were serum starved for 48 hours and proteins were extracted with lysis buffer (Sigma Cell Lytic Solution C2978) with 0.1% CHAPSO, 0.1% NP-40 and protease inhibitor (Complete EDTA-Free, Roche). Insoluble components were removed by centrifugation at 20000g. Flag beads (Anti Flag M2 Affinity Gel, Sigma A2250) were added to the cell extract and the mixture incubated for 2 hrs at 4°C. After centrifugation, beads were washed 3X with 0.1% Tween20/TBS and 3X with TBS before elution with 3XFLAG peptide (Sigma F4799). Gel loading buffer was added to the eluates for SDS-PAGE electrophoresis and Western blotting analysis.

Isolation of mRNA and quantitative mRNA analysis was performed as previously described (Jonassen et al., 2008) using the primers described in Table 3.

**Table 3.**
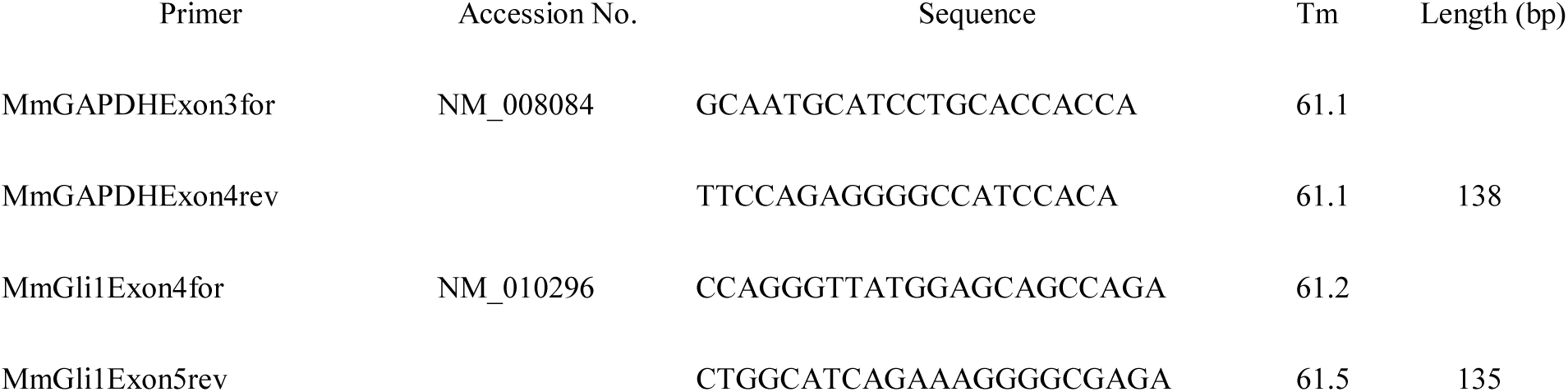
Quantitative RT-PCR Primers.

### Statistics

Data groups were analyzed by one-way ANOVA and compared using the Tukey posthoc test (GraphPad Prism, San Diego CA). Percentage data was arcsine transformed before analysis to normalize its distribution. Differences between groups were considered statistically significant if p < 0.05. Statistical significance is denoted with asterisks (*p=0.01 – 0.05; **p=0.01-0.001; ***p<0.001-0.0001, ****p<0.0001). Error bars are all S.D.

## Results

### Ubiquitinated Smo is retained in *Ift27^-/-^*, *Lztfl1^-/-^* and *Bbs2^-/-^* cilia but not in wild type cilia

Our previous work indicated that Ift25 and Ift27 are required for the regulated removal of Smo from cilia (Eguether et al., 2014; Keady et al., 2012). This suggests that hedgehog signaling regulates the interaction between Smo and the IFT particle, but the underlying mechanism is unknown. Structurally, Smo is a member of the G Protein-Coupled Receptor (GPCR) seven transmembrane protein family. Many GPCRs cycle between the cell surface and the endomembrane system depending upon ligand binding. The dynamics are often driven by beta-arrestin binding to activated receptor causing the receptor to associate with clathrin for internalization (Tian et al., 2014). Alternatively, the receptor may be ubiquitinated and internalized via the ESCRT machinery (Skieterska et al., 2017). While a published report indicates that Smo delivery to cilia required beta-arrestin (Kovacs et al., 2008), we were not able to observe any Smo trafficking defects in MEFs lacking beta-arrestin 1 and 2 (*Barr1/2*, Figure 1A,B).

**Figure 1.**
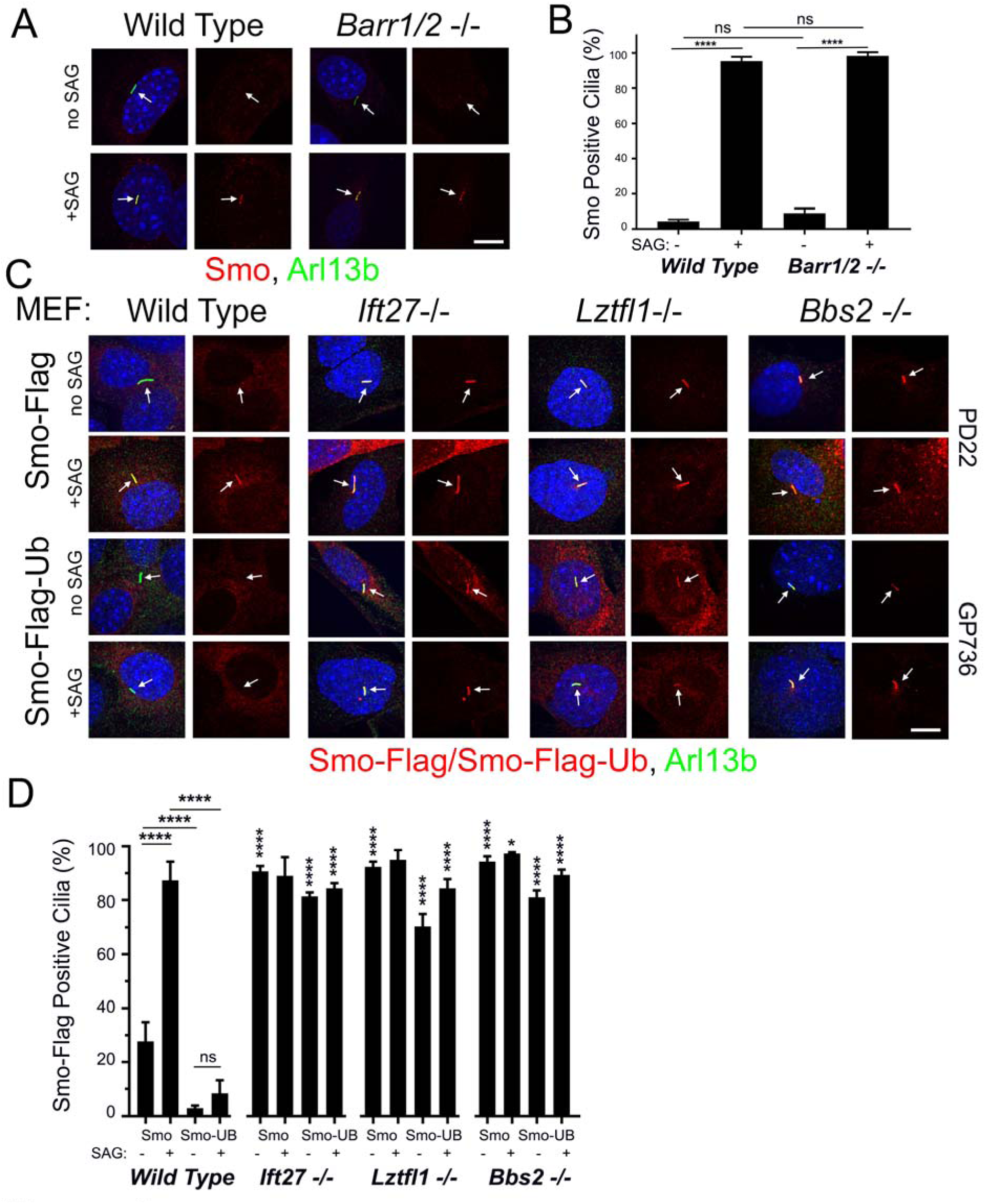
Smo-Ub Is Removed from Wild Type MEF Cilia but Not from *Ift27*, *Lztfl1* or *Bbs2* Mutant Cilia. **A.** Wild type and *Barr1/Barr2* double mutant MEFs were serum starved and stimulated with SAG for 24 hrs before being fixed and stained for endogenous Smo (red), cilia (Arl13b, green) and nuclei (4’,6-diamidino-2-phenylindole [DAPI], blue). Left image of each pair is a three-color composite, right image shows only the red Smo channel. Each image is maximum projection of a three-image stack taken at 0.2-micron intervals. Scale bar is 10 microns and applies to all images in panel. **B.** Presence of endogenous Smo or Smo-Ub in cilia was quantitated from the cells described in panel A. N is 3 replicates with at least 100 cells counted per condition. **** p<0.0001; ns, not significant. **C.** Wild type, *Ift27*, *Lztfl1* and *Bbs2* mutant MEFs were transfected with PD22 (Smo-Flag) or GP736 (Smo-Flag-Ub) and selected with Bsd. Confluent cells were serum starved and stimulated with SAG for 24 hrs before being fixed and stained for Smo or Smo-Ub (Flag, red), cilia (Arl13b, green) and nuclei (DAPI, blue). Left image of each pair is a three-color composite, right image shows only the red Flag channel. Each image is maximum projection of a three-image stack taken at 0.2-micron intervals. Scale bar is 10 microns and applies to all images in panel. **D.** Presence of Smo or Smo-Ub in cilia was quantitated from the cells described in panel C. N is 3 replicates with at least 100 cells counted per condition. **** p<0.0001; ns, not significant. For the mutants, significance is shown with respect to same condition in wild type.

To explore the role of ubiquitin in regulating the ciliary localization of Smo, we fused a single Ub open reading frame to the intracellular C-terminal end of Smo. This approach has been used to study the role of Ub in trafficking of GPCRs (Shih et al., 2000; Terrell et al., 1998). Cells expressing Smo-Flag from a CMV promoter show ∼25% Flag-positive cilia without pathway activation and the numbers are elevated to ∼90% positive cilia upon pathway activation by SAG. The addition of Ub to the C-terminus greatly reduces the amount of ciliary Smo under basal conditions and notably also blocks the accumulation usually associated with pathway activation by addition of SAG (Figure 1C,D). As we and others have previously observed for endogenous Smo (Eguether et al., 2014; Seo et al., 2011), Smo-Flag is retained in *Ift27^-/-^*, *Lztfl1^-/-^* and *Bbs2^-/-^* MEF cilia at both the basal state and after SAG induction. In contrast to the depletion of Smo-Flag-Ub from cilia that we saw in control cells, Smo-Flag-Ub is highly enriched in *Ift27^-/-^*, *Lztfl1^-/-^* and *Bbs2^-/-^*cilia (Figure 1C,D). Importantly, the expression of Smo-Flag-Ub does not interfere with the trafficking of endogenous Smo (Figure S1). These data suggest that the IFT/BBS system targets ubiquitinated Smo for ciliary removal.

To confirm our finding that Smo-Ub was retained in *Ift27^-/-^* mutant cilia but not in control cilia, we examined IMCD3 control and IMCD3*^Ift27-/-^* cells. In these kidney epithelial cells, Smo-Flag is retained in wild type cilia without pathway activation (Figure 2A,B). Confirming the MEF studies, Smo-Flag-Ub does not accumulate in control cilia but does accumulate in IMCD3*^Ift27-/-^* cilia (Figure 2A,B).

**Figure 2.**
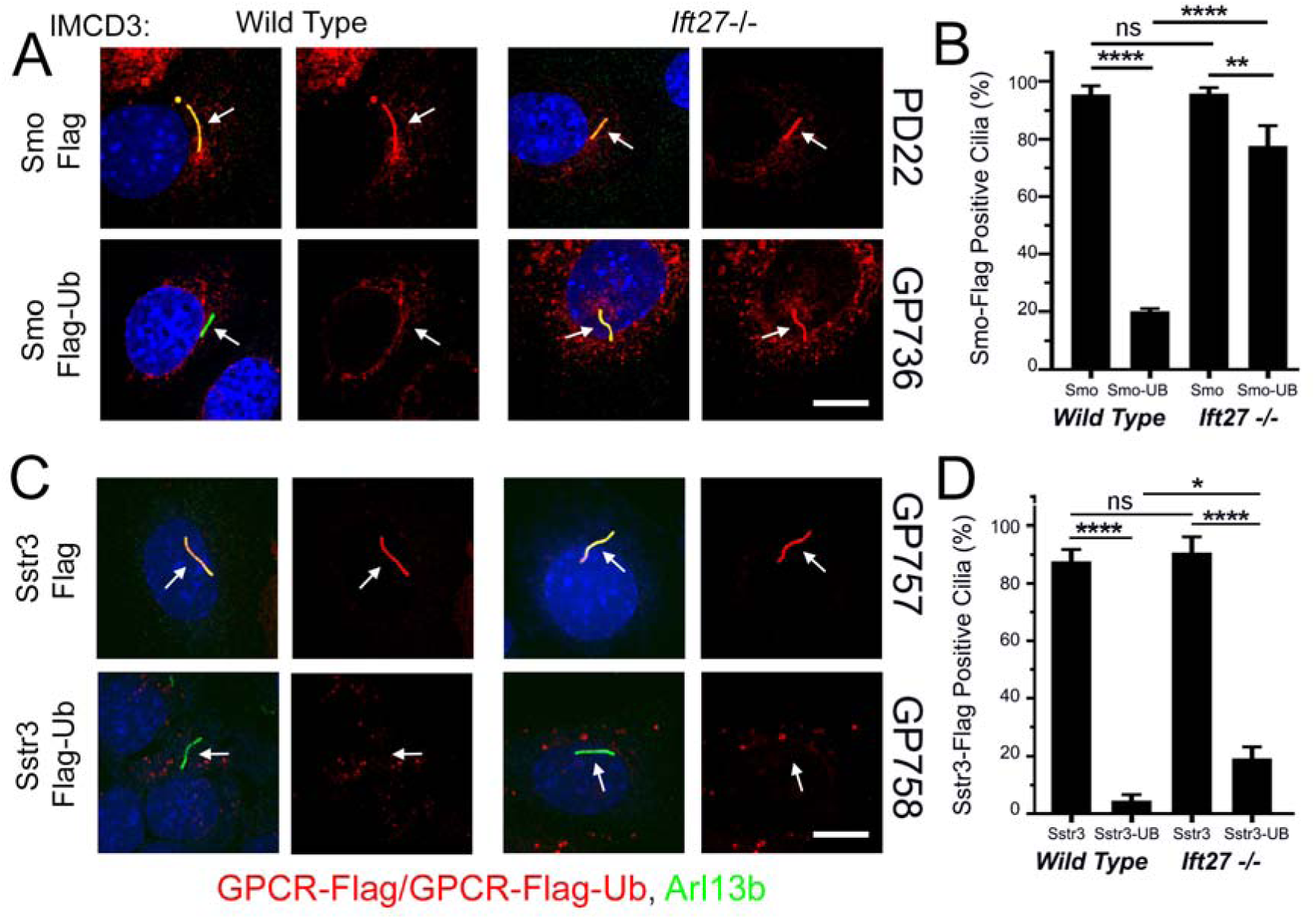
GPCR Trafficking in IMCD3 Cells Reveals Diversity in Ciliary Retrieval Mechanisms. **A.** Wild type and *Ift27* mutant IMCD3 cells were transfected with PD22 (Smo-Flag) or GP736 (Smo-Flag-Ub) and selected with Bsd. Confluent cells were serum starved for 48 hrs before being fixed and stained for Smo or Smo-Ub (Flag, red), cilia (Arl13b, green) and nuclei (DAPI, blue). Each image is maximum projection of a three-image stack taken at 0.2-micron intervals. Left image of each pair is a three-color composite, right image shows only the red Flag channel. Scale bar is 10 microns and applies to all images in the panel. **B.** Presence of Smo or Smo-Ub was quantitated from cells in panel A. N is 3 replicates with at least 100 cells counted per condition. **** p<0.0001; *** p<0.001; ns, not significant. **C.** Wild type and *Ift27* mutant IMCD3 cells were transfected with GP757 (Sstr3-Flag) or GP758 (Sstr3-Flag-Ub) and selected with Bsd. Confluent cells were serum starved for 48 hrs before being fixed and stained for Sstr3 or Sstr3-Ub (Flag, red), cilia (Arl13b, green) and nuclei (DAPI, blue). Each image is maximum projection of a three-image stack taken at 0.2-micron intervals. Left image of each pair is a three-color composite, right image shows only the red Flag channel. Scale bar is 10 microns and applies to all images in the panel. **D.** Presence of the Sstr3 or Sstr3-Ub was quantitated from cells in panel C. N is 3 replicates with at least 100 cells counted per condition. **** p<0.0001; ns, not significant.

To determine how other GPCRs behave, we tagged the somatostatin receptor Sstr3 with Flag and Ub. Sstr3-Flag is highly enriched in both control and IMCD3*^Ift27-/-^* cilia while Sstr3-Flag-Ub does not accumulate to significant levels in cilia of either cell line (Figure 2C,D). Similar results were obtained for the Ddr1 dopamine receptor (data not shown) suggesting that an Ift27-independent mechanism exists for the removal of some ubiquitinated cargos from cilia.

SmoM2 is an oncogenic form of Smo that contains a Trp to Leu mutation (MmSmo^W539L^) at cytoplasmic face of the last transmembrane domain (Xie et al., 1998). This mutation constitutively activates hedgehog signaling and SmoM2 is localized to cilia regardless of whether hedgehog ligand is present or not. Tagging SmoM2 with Ub removed it from wild type cilia as expected, but unexpectedly SmoM2-Ub was observed to be barely above background in *Ift27^-/-^*, *Lztfl1^-/-^* and *Bbs2^-/-^* MEF cilia (Figure 3A,B). Similar results were obtained in *Ift27^-/^*^-^ IMCD3 cells (Figure 3F,G). One possibility for the failure of SmoM2-Ub to accumulate in *Ift27^-/-^*, *Lztfl1^-/-^* and *Bbs2^-/-^* mutant cells is that perhaps SmoM2-Ub is highly unstable and degraded before it is able to traffic to cilia. To compare the relative stability of Smo-Ub and SmoM2-Ub, we treated cells expressing these proteins with MG132 for 4 hrs to build up levels of Smo-Ub or SmoM2-Ub, then added cycloheximide to prevent new synthesis and removed the MG132 to allow proteasomal degradation to proceed. MG132 treatment caused significant buildup of Smo-Ub and SmoM2-Ub in cells, which decayed with similar rates suggesting that it is not the stability of SmoM2-Ub that prevents its accumulation in *Ift27^-/-^* cilia (Supplemental Figure 3A). The large buildup of SmoM2-Ub caused by MG132 treatment was not sufficient to cause SmoM2-Ub to accumulate in *Ift27^-/-^* cilia (Figure 3C). Recently ectocytosis was proposed as an alternative pathway for clearing some GPCRs from cilia (Nager et al., 2017). This pathway is activated in *Bbs* mutants suggesting that perhaps ectocytosis can remove SmoM2-Ub from cilia on *Ift27^-/-^*, *Lztfl1^-/-^* and *Bbs2^-/-^* mutant cells. However, ectocytosis does not appear to be the explanation as SmoM2-Ub did not accumulate when ectocytosis was blocked with cytochalasin D (Figure 3C). Beta-arrestin-driven retrieval is another possible mechanism for removal of SmoM2-Ub, but this does not appear to be the mechanism as loss of *Barr1/2* did not cause accumulation of SmoM2-Ub in cilia (Figure 3C).

**Figure 3.**
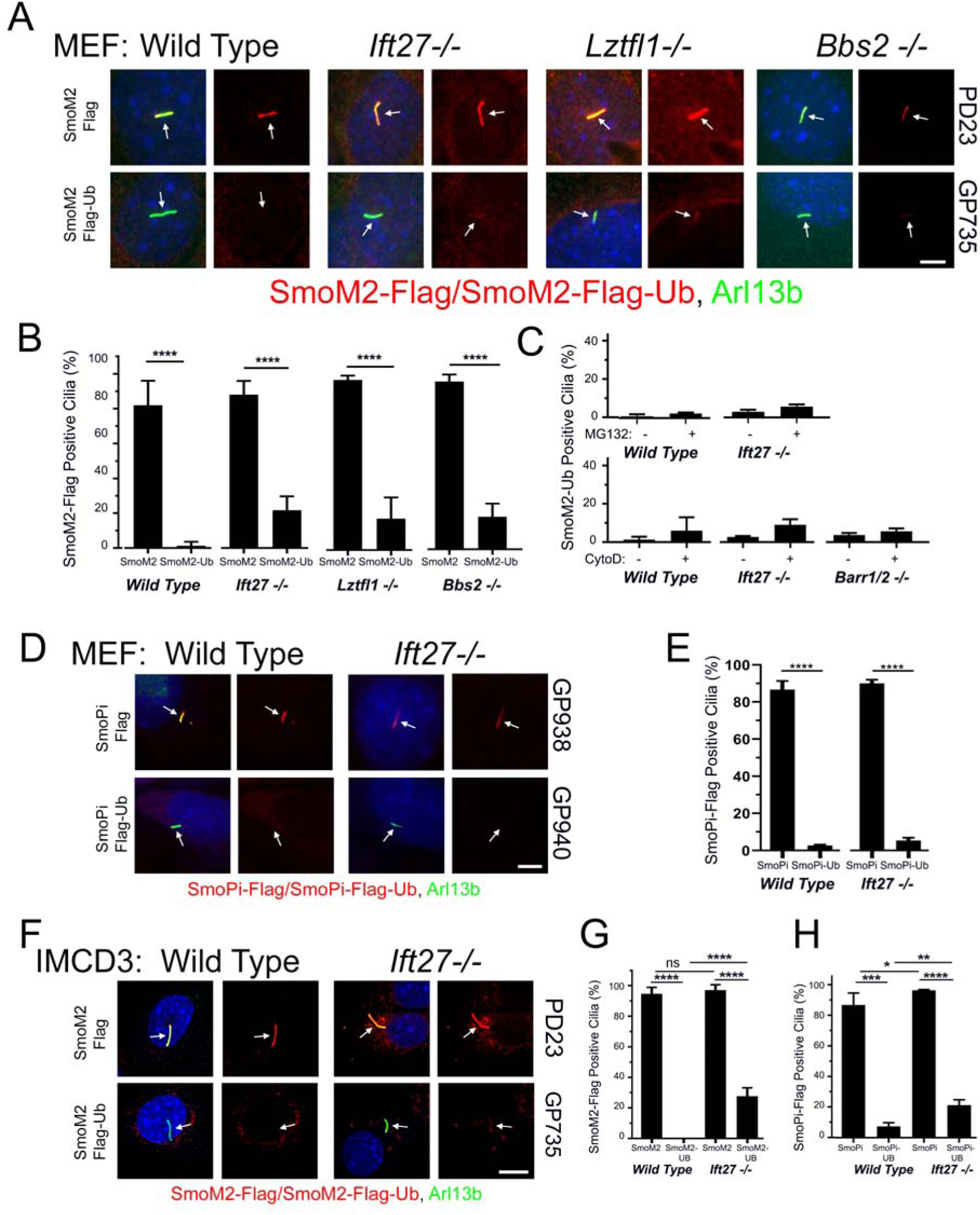
SmoM2 is different from Smo. A. Wild type, *Ift27*, *Lztfl1* and *Bbs2* mutant MEFs were transfected with PD23 (SmoM2-Flag) or GP735 (SmoM2-Flag-Ub) and selected with Bsd. Confluent cells were serum starved for 48 hrs before being fixed and stained for SmoM2 or SmoM2-Ub (Flag, red), cilia (Arl13b, green) and nuclei (DAPI, blue). Left image of each pair is a three-color composite, right image shows only the red Flag channel. Scale bar is 5 microns and applies to all images in panel. B. Presence of SmoM2 or SmoM2-Ub in cilia was quantitated from the cells described in panel A. N is 3 replicates with at least 100 cells counted per condition. **** p<0.0001. Mutants were not significantly different from same condition in wild type. C. Wild type, *Ift27* and *Barr1/2* mutant SmoM2-Ub cells treated like in panel B except that 1 micromolar MG132 for 4 hrs or 500 nanomolar cytochalasin D was added for 24 hrs before fixation. No differences were significant. D. Wild type and *Ift27* mutant MEFs were transfected with GP938 (SmoPi-Flag) or GP940 (SmoPi-Flag-Ub) and selected with Bsd. Confluent cells were serum starved for 48 hrs before being fixed and stained for SmoPi or SmoPi-Ub (Flag, red), cilia (Arl13b, green) and nuclei (DAPI, blue). Left image of each pair is a three-color composite, right image shows only the red Flag channel. Scale bar is 5 microns and applies to all images in panel. E. Presence of SmoPi or SmoPi-Ub in cilia was quantitated from the cells described in panel D. N is 3 replicates with at least 100 cells counted per condition. **** p<0.0001. *Ift27* mutants were not significantly different from same condition in wild type. F. Wild type and *Ift27* mutant IMCD3 cells were transfected with PD23 (SmoM2-Flag) or GP735 (SmoM2-Flag-Ub) and selected with Bsd. Confluent cells were serum starved for 48 hrs before being fixed and stained for Flag (red), cilia (Arl13b, green) and nuclei (DAPI, blue). Each image is maximum projection of a three-image stack taken at 0.2-micron intervals. Left image of each pair is a three-color composite, right image shows only the red Flag channel. Scale bar is 10 microns and applies to all images in panel. G. Presence of the SmoM2 or SmoM2-Ub was quantitated from cells in panels D. N is 3 replicates with at least 100 cells counted per condition. **** p<0.0001; *** p<0.001; ns, not significant. H. Presence of the SmoPi or SmoPi-Ub was quantitated from IMCD3 cells like in panel D except that they were transfected with SmoPi (GP938) and Smo-Pi-Ub (GP940). N is 3 replicates with at least 100 cells counted per condition. **** p<0.0001; *** p<0.001; ** p<0.01, * p<0.05, ns, not significant.

The observation that SmoM2 is retained in cilia in spite of the repressive activity of Ptch1 suggests that SmoM2 is in a conformation that is not a substrate for ubiquitination and removal by Ift27 and the BBSome. Huang et al. recently proposed a model for how the SmoM2 mutation shifts the structure of Smo towards the activated state. Their model proposes that in the inactive state, Trp539 at the end of helix 7 forms a Pi bond with Arg455 in helix 6. The SmoM2 mutation, which converts Trp539 to a Leu, would break this bond shifting the positions of transmembrane helixes 6 and 7 and the amphipathic helix that follows helix 7 (Supplemental Figure 3B)(Huang et al., 2018). SmoPi-Flag (Arg455Ala) mimicked SmoM2-Flag in that the receptor accumulated in cilia without the need for pathway activation and SmoPi-Flag-Ub was removed from cilia independent of Ift27 (Figure 3D,E). This finding suggests that Smo’s conformation influences the mechanism of removal from cilia with inactive Smo using an Ift27-dependent mechanism and activated SmoM2 and SmoPi using a retrieval pathway similar to the one used by Sstr3 and other GPCRs that are constitutively localized in cilia.

### Blocking ubiquitination causes Smo to accumulate in cilia

The ligation of ubiquitin to a substrate requires a cascade starting with an E1 ubiquitin-activating enzyme followed by an E2 ubiquitin-conjugating enzyme and lastly an E3 ubiquitin ligase. Pyr41 inhibits E1 enzymes (Yang et al., 2007) and we reasoned that if ubiquitination of Smo was required for its removal from cilia, Pyr41 should cause Smo to accumulate in cilia. While blocking an E1 activating enzyme is expected to affect many processes, we found that cells treated with 3.5 micromolar Pyr41 appeared healthy, remained ciliated and accumulated Smo in cilia. The Smo accumulation in cilia was detectable within 6 hrs of treatment and at 24 hrs was similar to the accumulation driven by SAG treatment (Figure 4A,C). Pyr41 did not appear to affect general IFT as Ift27 staining was similar in Pyr41-treated cells and controls (Figure 4B).

**Figure 4.**
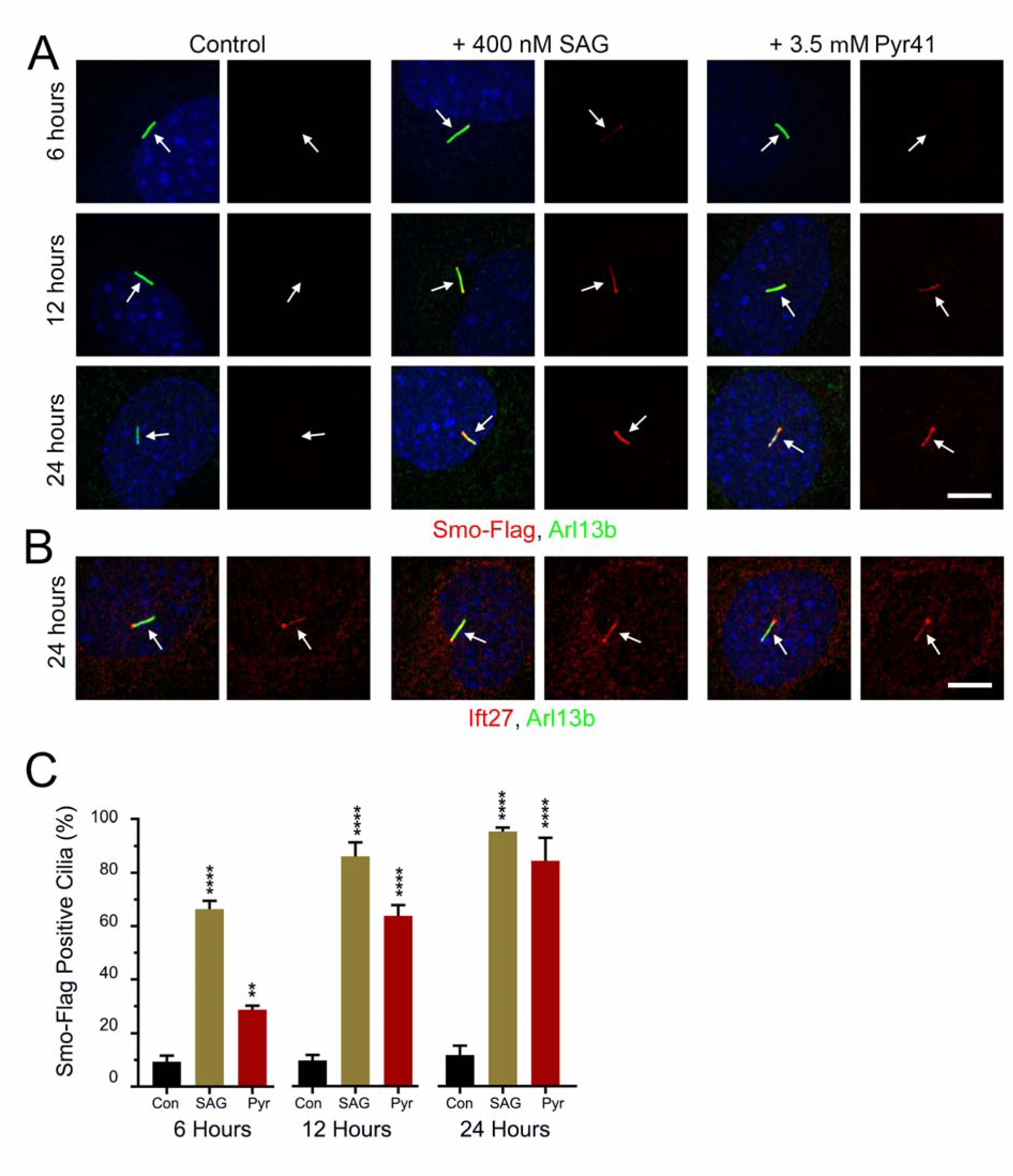
E1 inhibition Elevates Ciliary Smo Levels. **A.** Wild type MEFs were transfected with Smo-Flag (PD22) and a single-cell clone (11479.6T PD22 Clone3) was selected that showed low ciliary Smo-Flag at the basal level and high ciliary Smo-Flag after SAG stimulation. These cells were serum starved and not treated (negative control), treated with 400 nanomolar SAG (positive control) or treated with 3.5 micromolar Pyr41. Coverslips were removed at the indicated times, fixed and stained for Smo (Flag, red), cilia (Arl13b, green) and nuclei (DAPI, blue). Scale bar is 10 microns and applies to all images. Each image is maximum projection of a three-image stack taken at 0.2-micron intervals. Left image of each pair is a three-color composite, right image shows only the red Flag channel. **B.** Cells described in panel A were fixed at 24 hrs and stained for Ift27 (red), cilia (Arl13b, green) and nuclei (DAPI, blue). Scale bar is 10 microns and applies to all images. Each image is maximum projection of a three-image stack taken at 0.2-micron intervals. Left image of each pair is a three-color composite, right image shows only the red Ift27 channel. **C.** Presence of the Smo-Flag was quantitated from cells in panel A. N is 3 replicates with at least 100 cells counted per condition. ** p<0.01, **** p<0.0001 as compared to control at each time point.

### Lysines in Smo’s third intracellular loop are required for regulated ciliary localization

If blocking ubiquitination with Pyr41 is sufficient to cause Smo to accumulate in cilia, then mutating the site of ubiquitination should also cause Smo to accumulate. It is expected that a cytoplasmic lysine residue would be ubiquitinated and mouse Smo has 2 lysines in cytoplasmic loop two, 3 in cytoplasmic loop three and 16 in the C-terminal tail. Ubiquitination predictors gave inconsistent results without overlap of predicted sites of ubiquitination so Smo^noK^ was generated where all 21 cytoplasmic lysines were mutated to arginine (GP777). Smo^noK^ accumulated to high levels in cilia without pathway activation supporting the hypothesis that Ub is needed for ciliary removal (Figure 5A,B). To ensure that Smo^noK^ is still capable of being removed, we attached Ub and found that Smo^noK^-Ub was efficiently removed from the cilia. Ub^noK^ (all seven lysines of Ub are mutated to arginine) that is not a substrate for polyubiquitin was also effective in removing Smo^noK^ indicating that a single Ub is sufficient to remove Smo from the cilium (Figure 5A,B).

**Figure 5.**
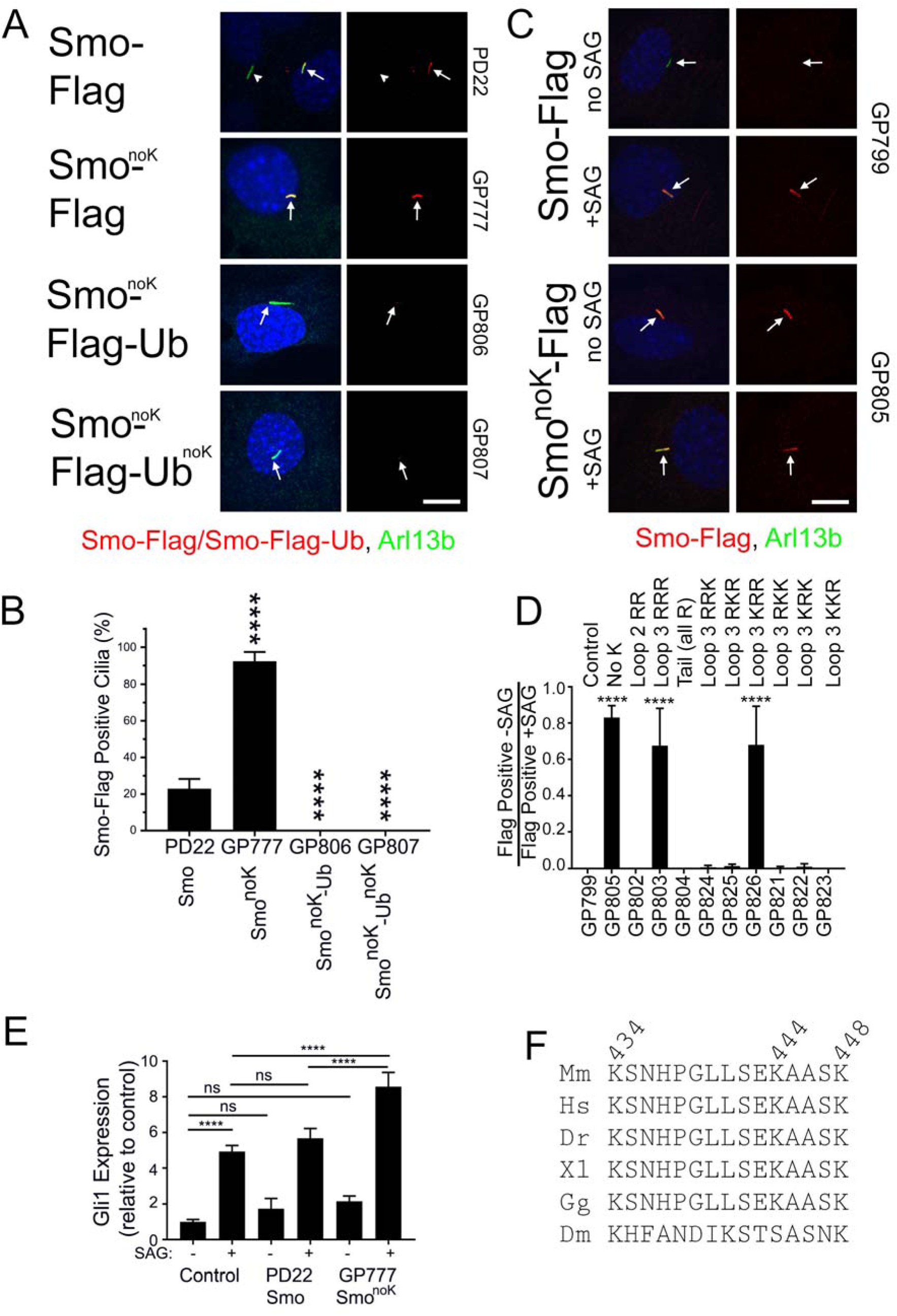
Smo Loop Three Lysines Are Required for Regulated Ciliary Localization. **A.** Wild type MEFs transfected with Smo-Flag (PD22), Smo^noK^-Flag (GP777), Smo^noK^-Flag-Ub (GP806) or Smo^noK^-Ub^noK^ (GP807) were serum starved, fixed and stained for Smo (Flag, red), cilia (Arl13b, green) and nuclei (DAPI, blue). Scale bar is 10 microns and applies to all images. Each image is maximum projection of a three-image stack taken at 0.2-micron intervals. Left image of each pair is a three-color composite, right image shows only the red Flag channel. **B.** Presence of the Smo-Flag or Smo-Flag-Ub was quantitated from cells in panel A. N is 3 replicates with at least 100 cells counted per condition. **** p<0.0001 as compared to control. **C.** Wild type MEFs transfected with GgCryD1-Bsd-IRES-Smo-Flag (GP799), GgCryD1-Bsd-IRES-Smo^noK^-Flag (GP805) were serum starved, treated +/-SAG, fixed and stained for Smo (Flag, red), cilia (Arl13b, green) and nuclei (DAPI, blue). Scale bar is 10 microns and applies to all images. Each image is maximum projection of a three-image stack taken at 0.2-micron intervals. Left image of each pair is a three-color composite, right image shows only the red Flag channel. **D.** Ciliary Smo was quantitated from the cells described in panel C as well as a series of mutants designed to identify the lysines important for regulated localization of Smo to cilia. Results are reported as a ratio of the percent of cells with Smo positive cilia without SAG divided by the percent of cells with Smo positive cilia after treatment with SAG. N is 3 replicates with at least 100 cells counted per condition. **** p<0.0001 compared to control (GP799). **E.** Wild type MEFs untransfected (control) or transfected with Smo-Flag (PD22) or Smo^noK^-Flag (GP777) were serum starved and treated +/- SAG for 24 hrs. RNA was collected, reverse transcribed and quantitated using real time PCR. Data are reported with respect to wild type unstimulated (-SAG). **** p<0.0001; ns, not significant. N=3 replicates. **F.** Sequence from Smo cytoplasmic loop 3 showing complete conservation in vertebrates (Mm, mouse; Hs, human; Dr, zebrafish; Xl, frog; Gg, chicken). *Drosophila* (Dm) retained three lysines but was otherwise divergent.

To understand the importance of Smo ubiquitination to signaling, we measured activation of the pathway (based on level of *Gli1* expression) in response to SAG in untransfected cells compared to ones transfected with Smo-Flag or Smo^noK^-Flag. Smo-Flag cells were similar to control cells, but Smo^noK^-Flag cells had a higher level of pathway activation suggesting that ubiquitination of Smo dampens the pathway (Figure 5E).

The observation that Smo^noK^ is retained in cilia suggested that we could identify the specific lysine residue(s) that is important for Smo removal by adding lysines back to Smo^noK^. This experiment is complicated by the fact that 20-25% of cilia were positive when wild type Smo was transfected into cells. We reasoned that this was due to the high level of expression of Smo driven by the CMV promoter saturating the ciliary removal system. To circumvent this problem, we created a construct with low Smo expression by replacing the CMV promoter with a weak GgCryD1 promoter, placing the Smo open reading frame downstream of an IRES and adding an out-of-frame methionine upstream of the Smo start codon (GP799). Cells expressing this construct did not show ciliary Smo until the pathway was activated (Figure 5C,D). However, drug selection did not work well for this construct, so the percent transfection was variable. To circumvent the problem of variable transfection, we ratioed the percent Smo-Flag positive without SAG treatment to the percent positive after SAG treatment (Figure 5D). Confirming the results with the CMV promoter (Figure 5A,B), Smo^noK^ expressed from the GgCryD1 promoter (GP805) accumulated in cilia without pathway activation (Figure 5C,D). Further analysis showed that the lysines in the C-terminal tail and loop 2 are not needed for removal of Smo-Flag from cilia but at least one of the three lysines in loop 3 (GP803) is critical. Further mutation of the loop 3 indicated that when both Lys444 and Lys448 are mutated to arginine (GP826), Smo is retained in cilia without pathway activation. Mutation of either of these lysines on their own was not enough to retain Smo (Figure 5D), suggesting that either residue can serve the regulatory role. This region of loop 3 is highly conserved in vertebrates with no substitutions in any of the main model organisms. In *Drosophila*, the region is not conserved except that it contains three lysines (Figure 5F).

### Smoothened ubiquitination is reduced by pathway activation

To explore the ubiquitination state of smoothened during pathway activation, stable cell lines expressing Smo-Flag and HA-Ub from different promoters were grown to confluence, serum starved for 24 hrs and then treated with or without hedgehog conditioned medium for an additional 24 hrs. After 20 hrs of hedgehog treatment, 1 micromolar MG132 was added for 4 hrs to block proteasomal degradation and then the cells were harvested for immunoprecipitation. Smoothened was precipitated with Flag antibody beads. Probing the immunoprecipitates for Flag showed that approximately equal amounts of smoothened were precipitated under both conditions (Figure 6A,B). Probing for HA showed a smear up the gel as expected for ubiquitinated proteins and showed that significantly more HA-Ub co-precipitated with Smo from control cells as compared to cells treated with SHH conditioned medium (Figure 6A,B). To ensure that the signal was due to ligation of Ub onto Smo, the experiment was repeated with Smo^noK^. As predicted, no HA-Ub was detected after immunoprecipitation of Smo^noK^ (Figure 6C,D). These data indicate that ubiquitination of Smo is controlled by hedgehog signaling.

**Figure 6.**
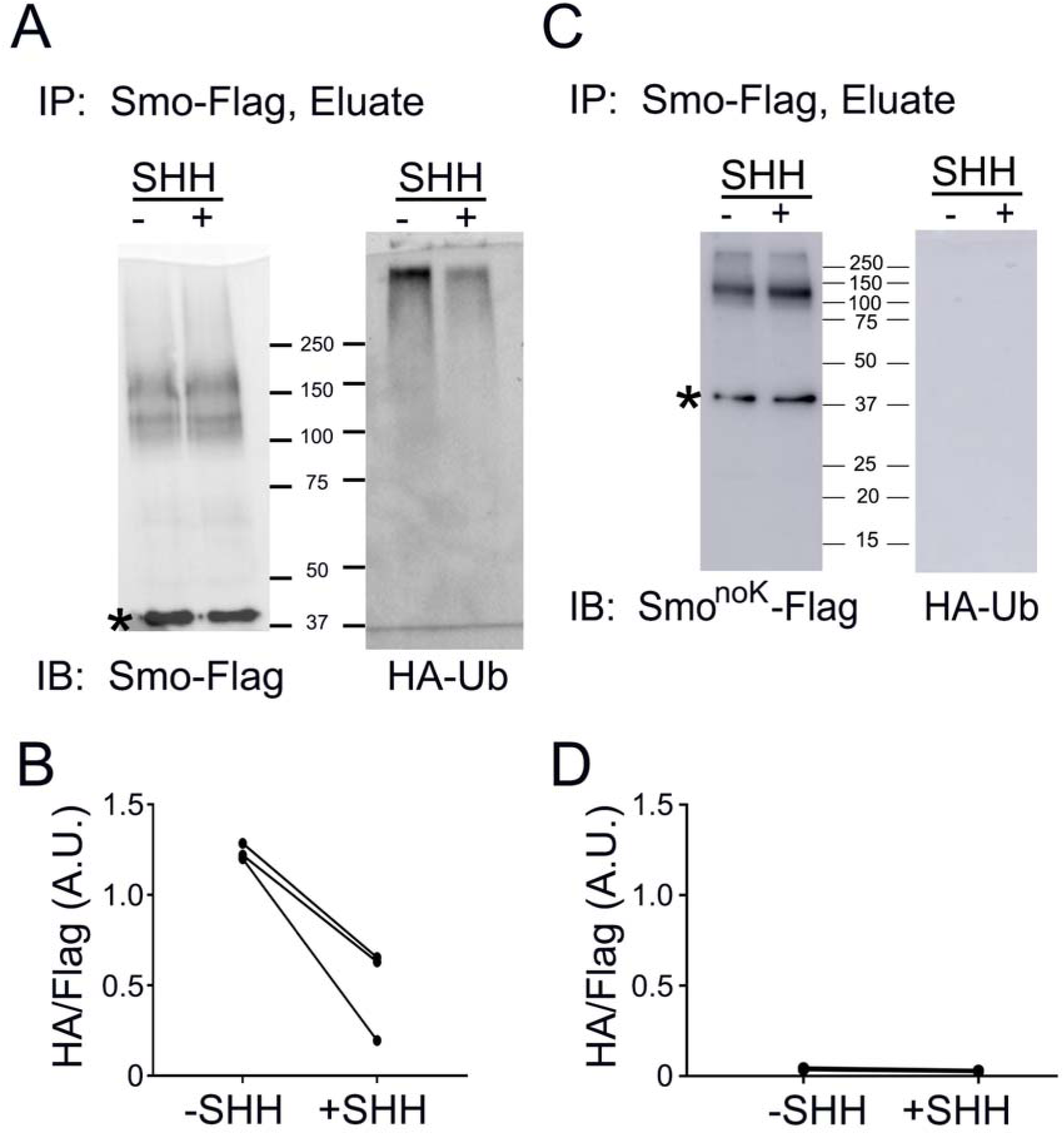
Ubiquitination of Smo Is Reduced by Pathway Activation. **A.** Wild type MEFs transfected with Smo-Flag (11479.6T+PD22_Clone3) and HA-Ub (PD14) were grown to confluence, serum starved 24 hrs, treated with or without hedgehog conditioned medium (SHH) for 24 hrs, treated with 1 micromolar MG132 for the last 4 hrs, and immunoprecipitated with anti-Flag beads. Precipitates were probed for Smo-Flag and HA-Ub. * marks an unrelated endogenous protein detected by the Flag antibody. **B.** Quantification of decreased Ub incorporation after treatment with hedgehog conditioned medium. N=3 repeats. P=0.03 with paired t-test. **C.** Wild type MEFs transfected with Smo^noK^-Flag (GP777) and HA-Ub (PD14) were grown to confluence, serum starved 24 hrs, treated with or without hedgehog conditioned medium (SHH) for 24 hrs, treated with 1micromolar MG132 for the last 4 hrs, and immunoprecipitated with anti-Flag beads. Precipitates were probed for Smo-Flag and HA-Ub. * marks an unrelated endogenous protein detected by the Flag antibody. **D.** Quantification of Ub incorporation after treatment with hedgehog conditioned medium. N=3 repeats. Differences are not significant.

## Discussion

During vertebrate hedgehog signaling, receptors and other components of the pathway are moved into and out of cilia. While these movements are critical to the proper regulation of the pathway, little is known about the molecular mechanisms driving them. In this work, we show that addition of Ub to the C-terminal end of Smo prevents its accumulation in wild type cilia in response to pathway activation. However, Smo-Ub still accumulates in cilia on cells lacking Ift27, Lztfl1 or Bbs2 suggesting that Ub couples Smo to the IFT system for removal from cilia. In support of this observation, blocking ubiquitination with an E1 inhibitor or by removing two critical lysines from Smo caused Smo to inappropriately accumulate in cilia in the off state. We further show that activation of hedgehog signaling reduces the amount of Ub on Smo. Taken together, these data suggest a model where an E3 ligase ubiquitinates Smo in the off state, which promotes its interaction with the IFT/BBSome for removal from cilia. Upon pathway activation, we expect this E3 ligase will be inactivated and there may also be a concomitant activation of a Smo deubiquitinase thus allowing Smo to remain in cilia (Figure 7).

**Figure 7.**
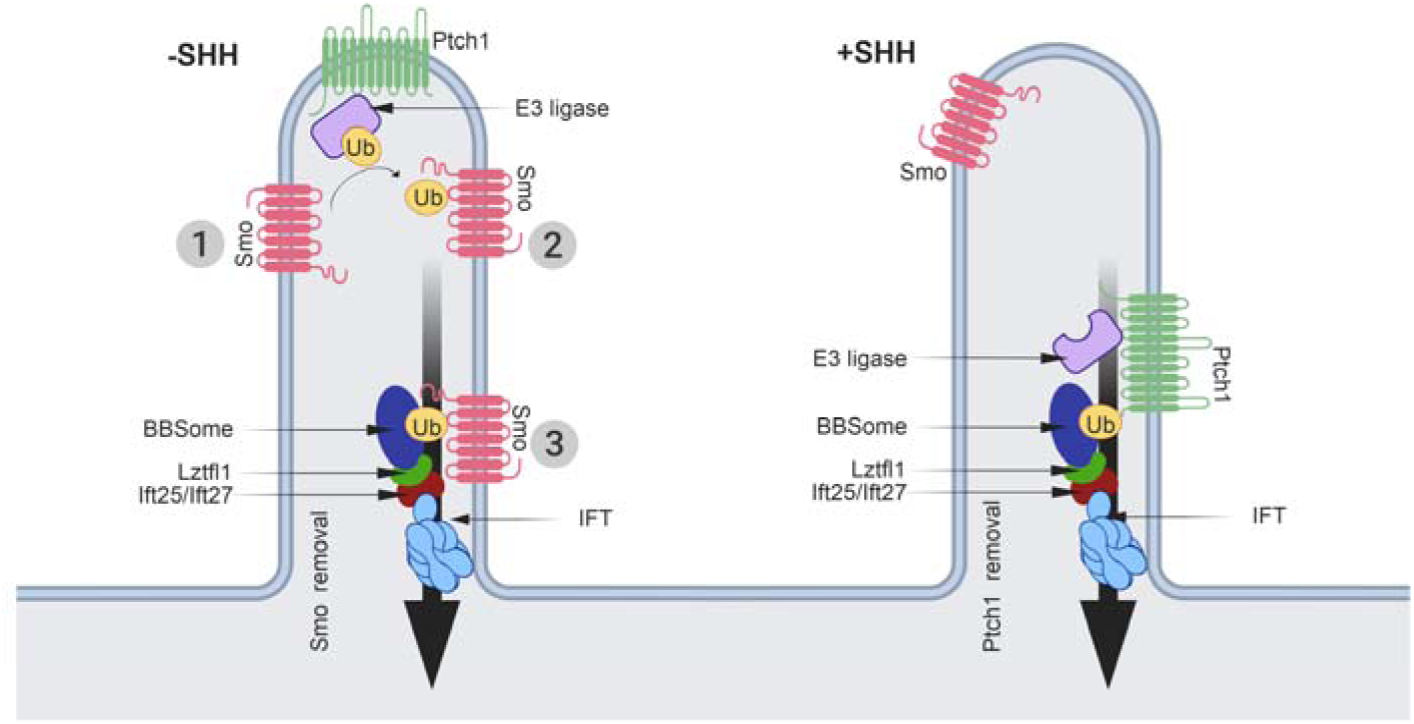
Model for the Mechanism Regulating Ciliary Levels of Smo. Our data and data from the literature supports a model where the ubiquitination state of Smo regulates its ciliary localization. In the simplest form of the model, Smo that enters cilia at the basal state (1) is ubiquitinated by unknown E3 ligase (2) on lysine residues in intracellular loop three, making it cargo for removal from the cilium by the IFT system (3). Activation of the pathway by SHH binding to Ptch1, would suppress the activity of the E3 ligase or if the E3 is ciliary localized, remove it from the cilium. This would allow Smo that enters the cilium to remain, become activated and trigger the downstream signaling events. A simple mechanism for regulating the ciliary localization of an E3 ligase could involve binding the ligase to Ptch1, which is known to bind E3 ligases. By attaching the ligase to Ptch1, it would be ciliary localized at the basal state and be removed upon pathway activation. While we have no data on the role of Ub in regulating Ptch1, a similar mechanism where Ptch1 is ubiquitinated by pathway activation could serve to keep ciliary Ptch1 levels low in the activated state.

The removal of ubiquitinated GPCRs from cilia appears to be general phenomena as the addition of Ub to Sstr3 prevented it from accumulating in cilia. However, in contrast to Smo-Ub, the removal of Sstr3-Ub was not dependent on Ift27. Unexpectedly, oncogenic SmoM2 behaved like Sstr3 rather than non-oncogenic Smo as SmoM2-Ub did not accumulate in cilia lacking Ift27. This suggests that an alternative mechanism exists to remove certain GPCRs from cilia. The removal was not blocked by loss of beta-arrestin or by inhibiting ectocytosis making it likely to be an IFT-dependent process. If it is IFT-dependent, then subunits of either IFT-A or IFT-B (besides Ift25 and Ift27) must be able to bind ubiquitinated GPCRs to remove them from cilia. Smo and SmoM2 differ by a single amino acid, Trp539Leu, located at the end of transmembrane 7. Trp539 is thought to interact with Arg455 to form a Pi lock that keeps Smo in the inactive configuration. Activation of Smo or mutating either Trp539 or Arg455 is thought to open the Pi lock, shifting transmembrane helixes 6 and 7 and causing amphipathic helix 8 in the C-terminal cytoplasmic tail to shift and partially unwind (Huang et al., 2018). Our finding that SmoPi-Ub is removed from Ift27 mutant cilia like SmoM2-Ub suggests that activation of Smo changes its conformation such that ubiquitinated forms no longer depend on the BBSome for removal. The detailed mechanism of how Smo interacts with the BBSome remains to be determined, but a short region just downstream of the SmoM2 mutation is reported to bind to BBS5 and BBS7 (Seo et al., 2011). This region contains the amphipathic helix that is shifted and partially unwound by the activation of Smo (Huang et al., 2018). It is possible that the SmoM2 and SmoPi mutations change the structure of this helix and the tail so it can be bound by other components of the IFT system for removal from cilia.

The role of Ub in ciliary biology and hedgehog signaling has not been extensively studied although several studies indicate that Ub is critical to both cilia and hedgehog signaling. In *Chlamydomonas*, Ub and isoforms of E1, E2 and E3 ligases are present in the cilium (Huang et al., 2009; Long et al., 2016; Pazour et al., 2005). Ciliary proteins become ubiquitinated while cilia are disassembling and ubiquitinated cargos accumulate in cilia when IFT is disrupted (Huang et al., 2009). In mammalian cells, VCP, which uses ATP to segregate Ub from binding partners is complexed with IFT-B though Ubxn10. Both VCP and Ubxn10 are required for ciliogenesis (Raman et al., 2015). In *Trypanosomes*, Ub co-purifies with the BBSome and BBSome mutations affect cellular localization of ubiquitinated receptors (Langousis et al., 2016). In *C. elegans*, attachment of Ub to the ciliary membrane protein Pkd2, reduced its level in cilia and similar to our findings, the BBSome appeared to be required (Hu et al., 2007; Xu et al., 2015).

Ub has been implicated in a number of steps of the hedgehog pathway ranging from the production of hedgehog ligand to regulating the proteolytic conversion of Gli3 full length to Gli3 repressor (Hsia et al., 2015). Less is known about the role of Ub in regulating the dynamics of Ptch1 and Smo although previous studies support a function for Ub in this process. In *Drosophila*, Smo is ubiquitinated in the off state and this drives sequestration in cytoplasmic vesicles (Ma et al., 2016; Xia et al., 2012; Zhou et al., 2018). The E3 ligase modifying *Drosophila* Smo is reported to be Herc4 (Jiang et al., 2019; Sun et al., 2019) and other studies suggest that the Usp8 and Uchl5 deubiquitinases regulate cellular localization of Smo. Herc4 was identified as a genetic modifier of hedgehog signaling and its loss stabilized Smo protein (Jiang et al., 2019; Sun et al., 2019). Usp8 was identified in an RNAi screen as a Smo deubiquitinase and its loss increased Smo ubiquitination (Xia et al., 2012). Uchl5 stabilizes *Drosophila* Smo and promotes its localization on the cell surface. The human homolog UCH37 was reported to promote Smo localization to cilia (Zhou et al., 2018). However, we used crispr to knock out Herc4 along with its close paralog Herc3, Usp8 and Uchl5/UCH37 and did not observe any effects on Smo localization suggesting that they are not the critical enzymes regulating Smo removal from mammalian cilia.

Ptch1 is also likely to be regulated by ubiquitination as the E3 ligases Smurf1 and Smurf2 ubiquitinate Ptch1 and promote its degradation (Yue et al., 2014). While this work focused on the degradation of Ptch1 by ubiquitination, it showed that knockdown of Smurf1 and Smurf2 increased the intensity of Ptch1-GFP in mammalian cilia (Yue et al., 2014). Another relevant study found two PY motifs, which are binding sites for HECT family ubiquitinases, in Ptch1 and showed that mutating these binding sites caused Ptch1 to remain in cilia after pathway activation (Kim et al., 2015). Proteomic analysis indicates that numerous E3 ligases bind the tail of Ptch1 (Yamaki et al., 2016).

Two recent large scale CRISPR screens of genes involved in hedgehog signaling identified a number of ubiquitin-related enzymes regulating hedgehog signaling (Breslow et al., 2018; Pusapati et al., 2018). Genes identified included E1, E2 and E3 ligases along with deubiquitinases and adaptor proteins. Pusapati and colleagues (Pusapati et al., 2018) identified three genes *Mgrn1*, *Megf8* and *Atthog* that all increased ciliary levels of Smo at the basal state. Of these, the loss *Megf8* and *Atthog* drove ciliary Smo to high levels that appear similar to what we observe with the loss of *Ift27* whereas *Mgrn1* loss was more variable with some cilia showing strong enrichment and others similar to controls. Currently it is not known if *Megf8* and *Atthog* are involved in ubiquitination whereas *Mgrn1* encodes an E3 ligase. This makes Mgrn1 a strong candidate to be the enzyme that regulates the ciliary localization of Smo but the variable amount of Smo accumulation in cilia suggests that there are likely other enzymes involved.

In summary, our work uncovers a mechanism for regulating the dynamic distribution of Smo during hedgehog signaling (Figure 7). In this model, hedgehog signaling regulates the activity or subcellular distribution of E3 ligases (and perhaps deubiquitinases) that modify Smo. At the basal state, the E3 ligase is active against Smo, perhaps by being localized to cilia. Any Smo that enters the cilium becomes ubiquitinated making it a substrate from removal by the IFT/BBSome. Upon activation of hedgehog signaling, the E3 ligase is inhibited and/or removed from the cilium. Thus, Smo that enters the cilium will not be ubiquitinated and will not interact with the IFT/BBS complex, instead it will remain in the cilium. The simplest mechanism for regulating the ciliary localization of an E3 ligase is to bind it to Ptch1, which is known to be removed from cilia upon pathway activation. As discussed previously, Ptch1 contains two PY motifs and is known to bind HECT family E3 ligases making this a plausible mechanism. In addition to regulating Smo distribution, ubiquitin could more generally regulate the ciliary levels of other receptors. The most likely being Ptch1. The removal of this receptor from cilia upon pathway activation could be driven by an E3 ligase activated by binding of ligand to Ptch1.

## Acknowledgments

We thank Dr. S. Doxsey (University of Massachusetts Medical School) for use of his microscope, Drs. K. Bellve, L. Lifshitz and K. Fogarty (University of Massachusetts Medical School Biomedical Imaging Group) for assistance with microscopy, Dr. W. Tsang (Institut de recherches cliniques de Montréal) for sharing HA-Ub and HA-Ub^noK^ constructs and Dr. A. Salic (Harvard Medical School) for sharing the SHH clone.

We thank Drs. J. Chen (Stony Brook University Medicine), Thibaut Eguether (Faculte de Medecine Pierre et Marie Curie), William Monis (Sanofi) and Abigail Smith (University of Massachusetts Medical School) for critical reading of this manuscript and for valuable input during its production.

This work was supported by the National Institutes of Health GM060992 and DK103632 to G.J.P.

---

**Figure S1. Related to Figure 1.**
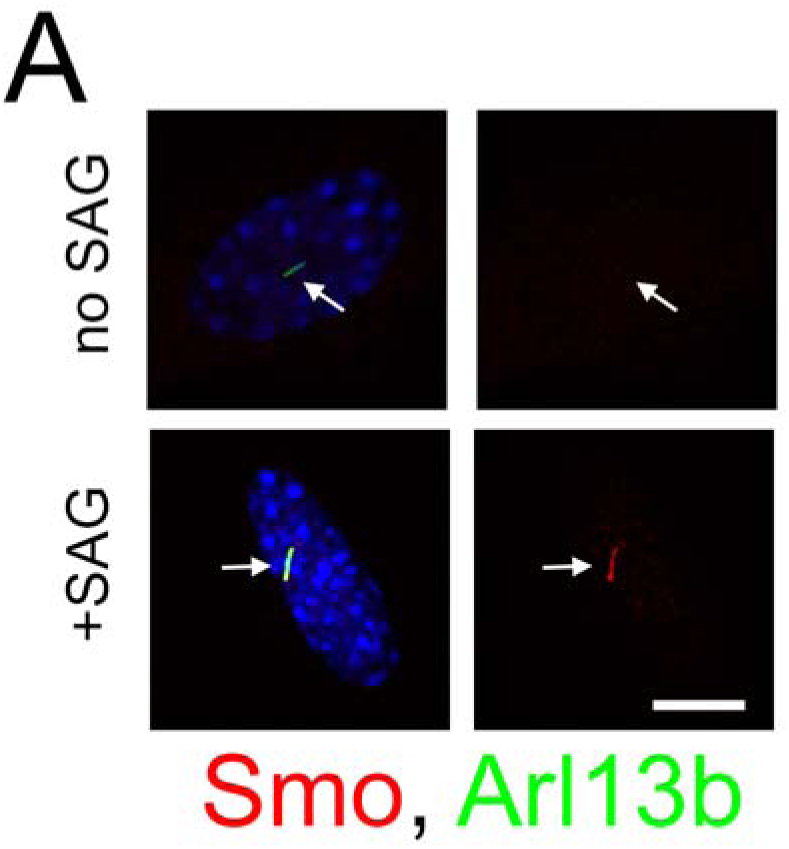
Endogenous Smo Is Trafficked Normally in Smo-Flag-Ub Expressing Cells. Wild type MEFs were transfected with GP736 (Smo-Flag-Ub) and selected with Bsd. Confluent cells were serum starved for 24 hrs and then stimulated with SAG for 24 hrs before being fixed and stained for endogenous Smo (red), cilia (Arl13b, green) and nuclei (4’,6-diamidino-2-phenylindole [DAPI], blue). Left image of each pair is a three-color composite, right image shows only the red Smo channel. Each image is maximum projection of a three-image stack taken at 0.2-micron intervals. Scale bar is 10 microns and applies to all images in panel. 3.67±3.2% of cells were Smo positive without SAG stimulation and 92.7±3.2% of cells were Smo positive after SAG stimulation (p<0.0001, Student’s t test).

**Figure S2. Related to Figure 1.**
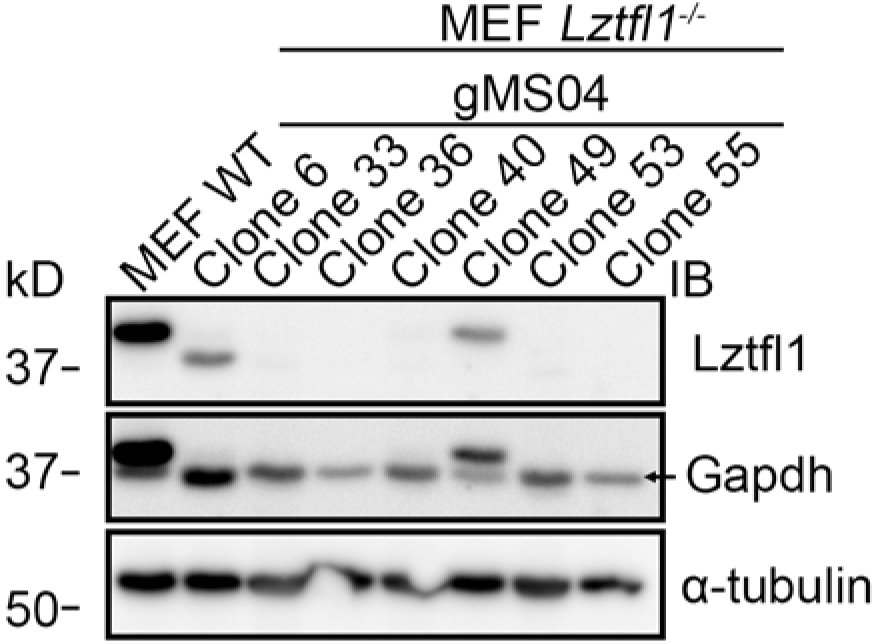
Genome Edited Cells. Western blot to document loss of Lztfl1 in edited MEFs. The Lztfl1 blot was reprobed with Gapdh and alpha tubulin antibodies to ensure relatively even loading. Note that the Lztfl1 bands in WT, clone 6 and clone 49 are visible in the Gapdh blot. Clone 36 was used in Figure 1.

**Figure S3. Related to Figure 3.**
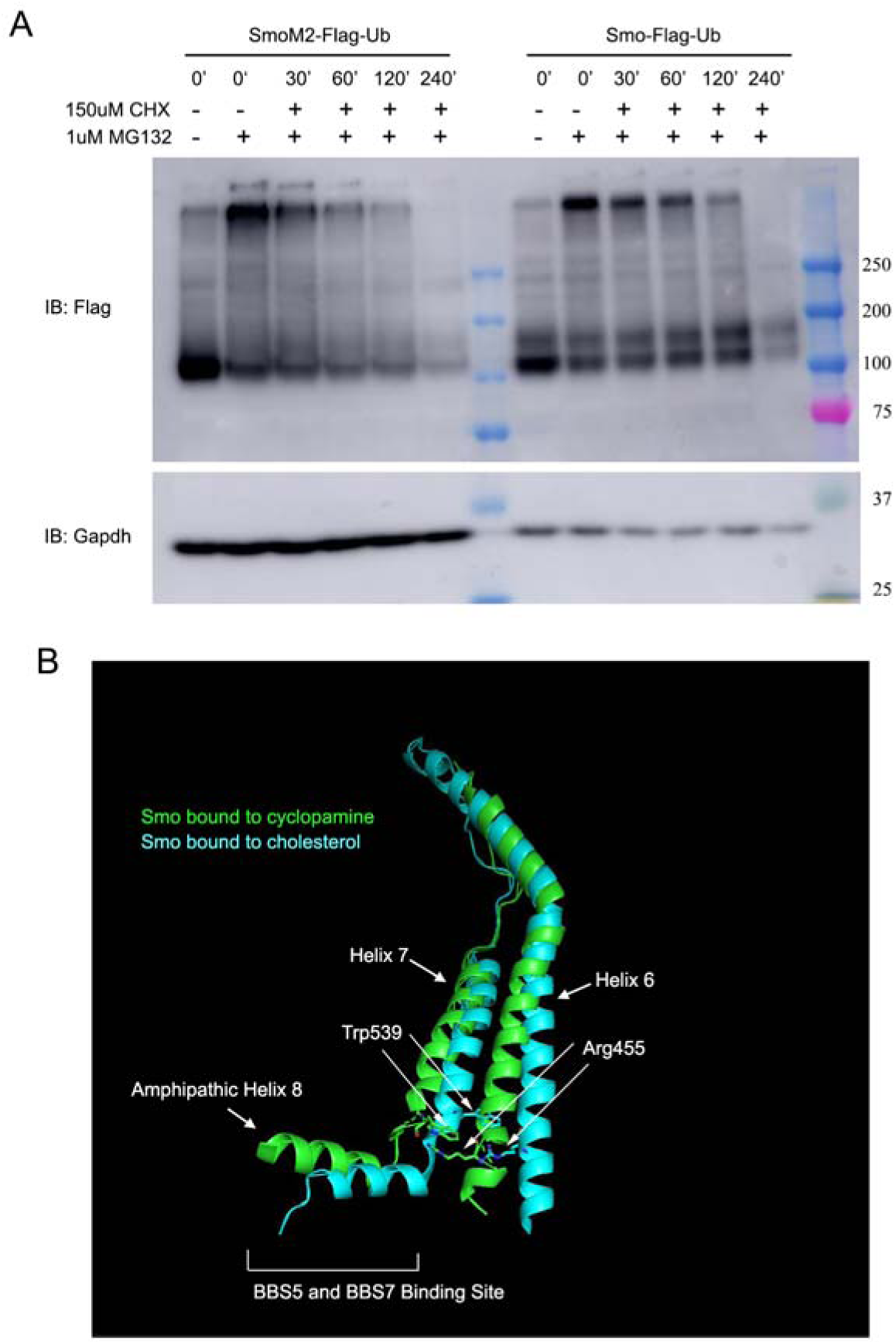
Structure of Smo. A. MEF cells expressing either SmoM2-Flag-Ub or Smo-Flag-Ub were treated with (+) or without (-) 1 micromolar MG132 for 4 hrs. After 4 hrs, 0’ time points were collected, the MG132 was washed out and 150 micromolar cycloheximide was added to the remaining cells. Samples were then collected at 30, 60, 120 and 240 min after the removal of MG132 and addition of cycloheximide. Half-life of Smo-Flag-Ub and SmoM2-Ub was 110 min and 181 min respectively (N=2). B. Ribbon diagram of Smo showing the location of the SmoM2 (W539) in transmembrane 7 and SmoPi (R455) residues in transmembrane 6 (redrawn by Pymol (PyMOL.org) from data in (Huang et al., 2018)). Blue green is cholesterol bound human Smo (5L7I) (Byrne et al., 2016) and green is cyclopamine-bound *Xenopus* Smo (6D32) (Huang et al., 2018). The two states predict the transitions from inactive (blue green) to active (green) when the Pi bond is broken. The breaking of the Pi bond is accompanied by shifts in the position of transmembrane helixes 6 and 7 and amphipathic helix 8, which is the proposed binding site of BBS5 and BBS7 (Seo et al., 2011).

## References

Breslow, D.K., S. Hoogendoorn, A.R. Kopp, D.W. Morgens, B.K. Vu, M.C. Kennedy, K. Han, A. Li, G.T. Hess, M.C. Bassik, J.K. Chen, and M.V. Nachury. 2018. A CRISPR-based screen for Hedgehog signaling provides insights into ciliary function and ciliopathies. Nat Genet. 50:460–471.

Byrne, E.F.X., R. Sircar, P.S. Miller, G. Hedger, G. Luchetti, S. Nachtergaele, M.D. Tully, L. Mydock-McGrane, D.F. Covey, R.P. Rambo, M.S.P. Sansom, S. Newstead, R. Rohatgi, and C. Siebold. 2016. Structural basis of Smoothened regulation by its extracellular domains. Nature. 535:517–522.

Corbit, K.C., P. Aanstad, V. Singla, A.R. Norman, D.Y. Stainier, and J.F. Reiter. 2005. Vertebrate Smoothened functions at the primary cilium. Nature. 437:1018–1021.

Eguether, T., F.P. Cordelieres, and G.J. Pazour. 2018. Intraflagellar transport is deeply integrated in hedgehog signaling. Mol Biol Cell. 29:1178–1189.

Eguether, T., J.T. San Agustin, B.T. Keady, J.A. Jonassen, Y. Liang, R. Francis, K. Tobita, C.A. Johnson, Z.A. Abdelhamed, C.W. Lo, and G.J. Pazour. 2014. IFT27 links the BBSome to IFT for maintenance of the ciliary signaling compartment. Dev Cell. 31:279–290.

Follit, J.A., R.A. Tuft, K.E. Fogarty, and G.J. Pazour. 2006. The intraflagellar transport protein IFT20 is associated with the Golgi complex and is required for cilia assembly. Mol Biol Cell. 17:3781–3792.

Haycraft, C.J., B. Banizs, Y. Aydin-Son, Q. Zhang, E.J. Michaud, and B.K. Yoder. 2005. Gli2 and Gli3 localize to cilia and require the intraflagellar transport protein polaris for processing and function. PLoS Genet. 1:e53.

Hsia, E.Y., Y. Gui, and X. Zheng. 2015. Regulation of Hedgehog signaling by ubiquitination. Front Biol (Beijing). 10:203–220.

Hu, J., S.G. Wittekind, and M.M. Barr. 2007. STAM and Hrs down-regulate ciliary TRP receptors. Mol Biol Cell. 18:3277–3289.

Huang, K., D.R. Diener, and J.L. Rosenbaum. 2009. The ubiquitin conjugation system is involved in the disassembly of cilia and flagella. J Cell Biol. 186:601–613.

Huang, P., S. Zheng, B.M. Wierbowski, Y. Kim, D. Nedelcu, L. Aravena, J. Liu, A.C. Kruse, and A. Salic. 2018. Structural Basis of Smoothened Activation in Hedgehog Signaling. Cell. 174:312–324 e316.

Huangfu, D., A. Liu, A.S. Rakeman, N.S. Murcia, L. Niswander, and K.V. Anderson. 2003. Hedgehog signalling in the mouse requires intraflagellar transport proteins. Nature. 426:83–87.

Jiang, W., X. Yao, Z. Shan, W. Li, Y. Gao, and Q. Zhang. 2019. E3 ligase Herc4 regulates Hedgehog signaling through promoting Smoothened degradation. J Mol Cell Biol.

Jonassen, J.A., J. San Agustin, J.A. Follit, and G.J. Pazour. 2008. Deletion of IFT20 in the mouse kidney causes misorientation of the mitotic spindle and cystic kidney disease. J Cell Biol. 183:377–384.

Keady, B.T., R. Samtani, K. Tobita, M. Tsuchya, J.T. San Agustin, J.A. Follit, J.A. Jonassen, R. Subramanian, C.W. Lo, and G.J. Pazour. 2012. IFT25 links the signal-dependent movement of Hedgehog components to intraflagellar transport. Dev Cell. 22:940–951.

Kim, J., E.Y. Hsia, A. Brigui, A. Plessis, P.A. Beachy, and X. Zheng. 2015. The role of ciliary trafficking in Hedgehog receptor signaling. Sci Signal. 8:ra55.

Kovacs, J.J., E.J. Whalen, R. Liu, K. Xiao, J. Kim, M. Chen, J. Wang, W. Chen, and R.J. Lefkowitz. 2008. Beta-arrestin-mediated localization of smoothened to the primary cilium. Science. 320:1777–1781.

Langousis, G., M.M. Shimogawa, E.A. Saada, A.A. Vashisht, R. Spreafico, A.R. Nager, W.D. Barshop, M.V. Nachury, J.A. Wohlschlegel, and K.L. Hill. 2016. Loss of the BBSome perturbs endocytic trafficking and disrupts virulence of Trypanosoma brucei. Proc Natl Acad Sci U S A. 113:632–637.

Liem, K.F., Jr., A. Ashe, M. He, P. Satir, J. Moran, D. Beier, C. Wicking, and K.V. Anderson. 2012. The IFT-A complex regulates Shh signaling through cilia structure and membrane protein trafficking. J Cell Biol. 197:789–800.

Long, H., F. Zhang, N. Xu, G. Liu, D.R. Diener, J.L. Rosenbaum, and K. Huang. 2016. Comparative Analysis of Ciliary Membranes and Ectosomes. Curr Biol. 26:3327–3335.

Ma, G., S. Li, Y. Han, S. Li, T. Yue, B. Wang, and J. Jiang. 2016. Regulation of Smoothened Trafficking and Hedgehog Signaling by the SUMO Pathway. Dev Cell. 39:438–451.

Nager, A.R., J.S. Goldstein, V. Herranz-Perez, D. Portran, F. Ye, J.M. Garcia-Verdugo, and M.V. Nachury. 2017. An Actin Network Dispatches Ciliary GPCRs into Extracellular Vesicles to Modulate Signaling. Cell. 168:252–263 e214.

Ocbina, P.J., and K.V. Anderson. 2008. Intraflagellar transport, cilia, and mammalian Hedgehog signaling: analysis in mouse embryonic fibroblasts. Dev Dyn. 237:2030–2038.

Pazour, G.J., N. Agrin, J. Leszyk, and G.B. Witman. 2005. Proteomic analysis of a eukaryotic cilium. J Cell Biol. 170:103–113.

Pazour, G.J., S.A. Baker, J.A. Deane, D.G. Cole, B.L. Dickert, J.L. Rosenbaum, G.B. Witman, and J.C. Besharse. 2002. The intraflagellar transport protein, IFT88, is essential for vertebrate photoreceptor assembly and maintenance. J Cell Biol. 157:103–113.

Pusapati, G.V., J.H. Kong, B.B. Patel, A. Krishnan, A. Sagner, M. Kinnebrew, J. Briscoe, L. Aravind, and R. Rohatgi. 2018. CRISPR Screens Uncover Genes that Regulate Target Cell Sensitivity to the Morphogen Sonic Hedgehog. Dev Cell. 44:113–129 e118.

Raman, M., M. Sergeev, M. Garnaas, J.R. Lydeard, E.L. Huttlin, W. Goessling, J.V. Shah, and J.W. Harper. 2015. Systematic proteomics of the VCP-UBXD adaptor network identifies a role for UBXN10 in regulating ciliogenesis. Nat Cell Biol. 17:1356–1369.

Rauchman, M.I., S.K. Nigam, E. Delpire, and S.R. Gullans. 1993. An osmotically tolerant inner medullary collecting duct cell line from an SV40 transgenic mouse. Am J Physiol. 265:F416–424.

Rohatgi, R., L. Milenkovic, and M.P. Scott. 2007. Patched1 regulates hedgehog signaling at the primary cilium. Science. 317:372–376.

Sanjana, N.E., O. Shalem, and F. Zhang. 2014. Improved vectors and genome-wide libraries for CRISPR screening. Nat Methods. 11:783–784.

Seo, S., Q. Zhang, K. Bugge, D.K. Breslow, C.C. Searby, M.V. Nachury, and V.C. Sheffield. 2011. A novel protein LZTFL1 regulates ciliary trafficking of the BBSome and Smoothened. PLoS Genet. 7:e1002358.

Shih, S.C., K.E. Sloper-Mould, and L. Hicke. 2000. Monoubiquitin carries a novel internalization signal that is appended to activated receptors. EMBO J. 19:187–198.

Skieterska, K., P. Rondou, and K. Van Craenenbroeck. 2017. Regulation of G Protein-Coupled Receptors by Ubiquitination. Int J Mol Sci. 18.

Sun, X., B. Sun, M. Cui, and Z. Zhou. 2019. HERC4 exerts an anti-tumor role through destabilizing the oncoprotein Smo. Biochem Biophys Res Commun.

Swatek, K.N., and D. Komander. 2016. Ubiquitin modifications. Cell Res. 26:399–422.

Terrell, J., S. Shih, R. Dunn, and L. Hicke. 1998. A function for monoubiquitination in the internalization of a G protein-coupled receptor. Mol Cell. 1:193–202.

Tian, X., D.S. Kang, and J.L. Benovic. 2014. beta-arrestins and G protein-coupled receptor trafficking. Handb Exp Pharmacol. 219:173–186.

Wang, Q., Z. Peng, H. Long, X. Deng, and K. Huang. 2019. Polyubiquitylation of alpha-tubulin at K304 is required for flagellar disassembly in Chlamydomonas. J Cell Sci. 132.

Wilson, A.A., L.W. Kwok, A.H. Hovav, S.J. Ohle, F.F. Little, A. Fine, and D.N. Kotton. 2008. Sustained expression of alpha1-antitrypsin after transplantation of manipulated hematopoietic stem cells. Am J Respir Cell Mol Biol. 39:133–141.

Xia, R., H. Jia, J. Fan, Y. Liu, and J. Jia. 2012. USP8 promotes smoothened signaling by preventing its ubiquitination and changing its subcellular localization. PLoS Biol. 10:e1001238.

Xie, J., M. Murone, S.M. Luoh, A. Ryan, Q. Gu, C. Zhang, J.M. Bonifas, C.W. Lam, M. Hynes, A. Goddard, A. Rosenthal, E.H. Epstein, Jr., and F.J. de Sauvage. 1998. Activating Smoothened mutations in sporadic basal-cell carcinoma. Nature. 391:90–92.

Xu, Q., Y. Zhang, Q. Wei, Y. Huang, Y. Li, K. Ling, and J. Hu. 2015. BBS4 and BBS5 show functional redundancy in the BBSome to regulate the degradative sorting of ciliary sensory receptors. Sci Rep. 5:11855.

Yamaki, Y., H. Kagawa, T. Hatta, T. Natsume, and H. Kawahara. 2016. The C-terminal cytoplasmic tail of hedgehog receptor Patched1 is a platform for E3 ubiquitin ligase complexes. Mol Cell Biochem. 414:1–12.

Yang, Y., J. Kitagaki, R.M. Dai, Y.C. Tsai, K.L. Lorick, R.L. Ludwig, S.A. Pierre, J.P. Jensen, I.V. Davydov, P. Oberoi, C.C. Li, J.H. Kenten, J.A. Beutler, K.H. Vousden, and A.M. Weissman. 2007. Inhibitors of ubiquitin-activating enzyme (E1), a new class of potential cancer therapeutics. Cancer Res. 67:9472–9481.

Yue, S., L.Y. Tang, Y. Tang, Y. Tang, Q.H. Shen, J. Ding, Y. Chen, Z. Zhang, T.T. Yu, Y.E. Zhang, and S.Y. Cheng. 2014. Requirement of Smurf-mediated endocytosis of Patched1 in sonic hedgehog signal reception. Elife. 3.

Zhou, Z., X. Yao, S. Pang, P. Chen, W. Jiang, Z. Shan, and Q. Zhang. 2018. The deubiquitinase UCHL5/UCH37 positively regulates Hedgehog signaling by deubiquitinating Smoothened. J Mol Cell Biol. 10:243–257.

